# Predictive processing of scene layout depends on naturalistic depth of field

**DOI:** 10.1101/2021.11.08.467670

**Authors:** Marco Gandolfo, Hendrik Nägele, Marius V. Peelen

**Affiliations:** Donders Institute for brain, cognition, and behaviour, Radboud University, 6525HR, Nijmegen, NL

**Keywords:** Visual perception, Predictive processing, Scene memory, Natural vision, Depth of field, Visual memory

## Abstract

Boundary extension (BE) is a classic memory illusion in which observers remember more of a scene than was presented. According to predictive processing accounts, BE reflects the integration of visual input and expectations of what is beyond a scene’s boundaries. According to normalization accounts, BE rather reflects one end of a normalization process towards a scene’s typically-experienced viewing distance, such that close-up views give BE but distant views give boundary contraction. Here across four experiments, we show that BE strongly depends on depth-of-field (DOF), as determined by the aperture settings on a camera. Photographs with naturalistic DOF led to larger BE than photographs with unnaturalistic DOF, even when showing distant views. We propose that BE reflects a predictive mechanism with adaptive value that is strongest for naturalistic views of scenes. The current findings indicate that DOF is an important variable to consider in the study of scene perception and memory.

**Statement of Relevance:** In daily life, we experience a rich and continuous visual world in spite of the capacity limits of the visual system. We may compensate for such limits with our memory, by filling-in the visual input with anticipatory representations of upcoming views. The boundary extension illusion (BE) provides a tool to investigate this phenomenon. For example, not all images equally lead to BE. In this set of studies, we show that memory extrapolation beyond scene boundaries is strongest for images resembling human visual experience, showing depth-of-field in the range of human vision. Based on these findings, we propose that predicting upcoming views is conditional to a scene being perceived as naturalistic. More generally, the strong reliance of a cognitive effect, such as BE, on naturalistic image properties indicates that it is imperative to use image sets that are ecologically-representative when studying the cognitive, computational, and neural mechanisms of scene processing.

## Introduction

Boundary extension (BE) is a classic memory illusion in which observers remember more of a scene than was actually presented (Intraub et al., 1989). For example, when participants draw a previously observed scene, they typically include information not present in the photograph, expanding the scene beyond its boundaries (Intraub et al., 1989). Similarly, when rating the viewpoint of a second probe photograph relative to a remembered target photograph, participants rate the exact same view as closer than the original image, indicating that they remembered more of the scene than was presented (Bertamini et al., 2005; Intraub & Dickinson, 2008; Dickinson and Intraub, 2008, Bainbridge & Baker, 2020).

Why does this illusion occur? One possibility is that it reflects a predictive mechanism, in which scene percepts do not correspond solely to the visual input but are rather *constructed* through integration of visual input and expectations of what is beyond a scene’s boundaries (Intraub, 2010; Maguire & Mullally, 2013). The discrete views sampled by the human eye may thus be supplemented with anticipatory representations of the scene, providing a continuous visual experience.

Alternatively, BE may reflect one end of a generic normalization process in memory (Bartlett, 1932), where the perceived scene viewpoint regresses towards a prototypical viewing distance: A view that is close-up compared to the observer’s average viewing distance of that scene will be remembered as farther away (yielding BE), whereas far-away views will be remembered as closer-up, generating the opposite error, boundary contraction (BC). On this account, BE and BC are in principle equally common, only depending on whether the presented photographs are taken from a relatively close or far distance.

Previous work has shown that BE occurs more frequently than BC (Hubbard et al., 2010; Intraub, 2020). Furthermore, BE occurs for scenes but not for single, isolated objects (Gottesman & Intraub, 2002; Intraub et al., 1998), unlike normalization, which also occurs for isolated objects (Brady, et al., 2011; Konkle & Oliva, 2007; Intraub, et al., 1998). However, while these results appear to show that BE cannot be entirely accounted for by normalization, this conclusion relies on the strong assumption that the scene photographs used in previous work were representative of our daily-life visual experience, particularly with respect to viewing distance. If the photographs used in BE studies were not representative, for example by primarily showing views that were closer than usually experienced, BE may be entirely explained by a normalization account.

Indeed, using a large and diverse stimulus set (1000 photographs), BE was recently shown to be equally common as BC (Bainbridge & Baker, 2020). Moreover, the direction of these memory errors was correlated with subjective image distance ratings: in line with the normalization account, subjectively close scenes tended to extend, and subjectively far scenes tended to contract. These results suggest that BE may be one side of a regression towards the typical scene-viewing distance.

The above discussion highlights that approximating daily-life visual experience is critical for both accounts: for the predictive processing account because it proposes that BE reflects expectations based on real-world experience; and for the normalization account because normalization is relative to our prior experience. If BE reflects expectations, it should be observed more prominently (and be more common than BC) when using views representative of our visual experience. Importantly, this is not automatically achieved through the use of large and diverse image sets. Even if these are representative of pictorial views of scenes (that is, of how scene photographs are taken), they are not necessarily representative of how an observer perceives their surroundings during day-to-day visual experience.

A striking example is that a camera can produce pictures with a deeper depth-of-field (DOF) than the eye can (Middleton, 1958; Artal, 2014). The DOF represents the distance range over which an object may be moved without causing a sharpness reduction. On a camera, DOF can be controlled by the lens aperture, typically specified by a f-number (focal length divided by aperture diameter). The eye’s DOF similarly depends on the aperture of the pupil. The larger the aperture, the shallower the depth of field, resulting in a smaller distance range within which objects are in focus. The aperture range of most cameras is much larger than that of the eye, which ranges between f/2.1 and f/8. Specifically, photographs often have very deep DOF (e.g., f/22), such that everything in the picture is sharp, leading to views of scenes that are highly unnatural.

Here, we show that BE strongly depends on DOF, as determined by the aperture settings on a camera. Re-analyzing the 1000-image dataset (Bainbridge & Baker, 2020), we find that BE was much more common than BC for photographs with DOFs within the range of human vision. Subsequently, in three experiments, we demonstrate the specific influence of DOF on BE while controlling for factors that may naturally co-vary with DOF in scene photographs, including viewing distance, number and type of objects, and image statistics. Again, BE was strongest for the relatively shallow DOF that characterizes human vision, and was observed even for distant views of scenes. Altogether, our results are most in line with views of BE as a predictive mechanism, with this error being reliably observed for naturalistic views of scenes to which the brain has adapted.

### Experiment 1

The goal of Experiment 1 was to test for a relationship between the perceived DOF of a photograph and its boundary transformation scores (BE or BC) in the large image set recently used by Bainbridge & Baker (2020). To this end, we collected aperture ratings (as judged from DOF) for each of the 1000 scenes and related these ratings to the boundary transformation scores made available by the original study. Based on the predictive processing account, we hypothesized that the more an image would be rated to be shot with large aperture – resembling natural vision – the more strongly it would elicit boundary extension.

## Methods

### Participants

Twenty-six participants (22 females, mean age = 24.5 ± 5.84) signed up for the online scene rating experiment. Participants were recruited through the participant panel of the University (SONA) in return for course credits and through social media. The experiment was advertised for people who knew the basics of photography. To further ensure participants’ expertise, prior to the scene ratings, participants went through a photography quiz. First, participants were asked to define their photography experience with a multiple-choice question (i.e., How would you define your experience with photography? 1) Only took pictures with my smartphone; 2) Some knowledge, but no hands-on experience with Single-Lens-Reflex (SLR) cameras; 3) Some experience using SLR cameras; 4) (Semi-) Professional photographer). Second, participants had to complete a quiz with 5 multiple-choice questions. Four of these questions asked how a photograph would look like if shot with large aperture (i.e., shallow depth of field, lots of light entering the camera) and small aperture (i.e., deep depth of field, little light entering the camera). Finally, one question asked whether a lens aperture of f/5.6 was larger or smaller than f/22.0, testing their knowledge of f-stop values. Based on the above questions, 6 participants were excluded because they signed up and then declared to have no experience with single lens reflex photography (3 participants) or because they failed 3 or more questions out of 5 in the photography quiz (4 errors for 2 participants, 5 errors for 1 participant). Among the 20 included participants, only 5 participants committed one or two errors (2 participants – 1 error; 3 participants – 2 errors). Quiz data for all the participants are available in the OSF repository. This and the following studies were approved by the Faculty’s Ethics Committee (ECSW2017-2306).

### Stimuli and apparatus

Stimuli were taken from Bainbridge and Baker (2020), corresponding to the 1000 images of their rapid serial visual presentation (RSVP) experiments. These were downloaded from the OSF link provided in their paper. Further, for the practice trials, 24 images were downloaded from Shutterdial (shutterdial – search images by camera settings). This website allows to search for photos according to their metadata (including aperture) from the Flickr API. We selected twelve pictures with large aperture *f* <= 4 and twelve pictures with small aperture *f* >= 22.0 so that the difference between shallow and deep depth of field was salient. This and the following experiments were online, coded in Javascript using the JSPsych toolbox 6.2.0 (de Leeuw, 2015). The data were saved on the Pavlovia.org servers (www.pavlovia.org).

### Procedure

Following the photography quiz, participants were shown the task instructions. Here, the concept of aperture was briefly re-explained. Afterwards, each participant was presented with 6 randomly selected practice images (3 with large, and 3 with small aperture). In the practice, participants simply had to report whether an image was shot with large or small aperture using the mouse. They received feedback after each trial. When the practice was completed, more detailed example images were presented, this time stressing the continuity of the aperture effect over the depth of field of the pictures. This was achieved by showing images of the same photograph shot at different levels of aperture (i.e., large – middle – small). Finally, participants were informed that the upcoming ratings would be performed on a slider from 0 (large) to 100 (small) aperture. Each rating trial began with a fixation cross with a random duration of 1, 1.25 or 1.5 seconds. Following fixation, participants saw an image appear on screen concurrently with the slider positioned in the middle (aperture = 50). To continue to the next trial, participants had to touch the slider on the scale and press the button “continue” placed below the image and the slider. After every 50 ratings, participants were invited to take a break.

### Design

Each participant completed 1 of 4 versions of the experiment. Each version included 250 images from Bainbridge & Baker (2020), equally divided between the SUN database (Xiao et al., 2016) and the Google Open Images (Kuznetsova et al., 2020) database, where the images were originally from. The images were presented in random order to each participant. In total, we obtained 5 aperture ratings for each of the 1000 stimuli of Bainbridge & Baker (2020), matching the number of subjective distance ratings collected by Bainbridge & Baker (2020).

### Analysis

All the analyses were conducted using R (R core team, version 4.0.1). We calculated the mean aperture rating for each of the 1000 images. We also split these ratings (from 0-100) in 5 categorical bins of 20 (XL 0-20; L 21-40; M 41-60; S 61-80; XS 81-100). Afterwards, we downloaded the raw data coming from the 1000-image RSVP Experiments 1 and 2 (Bainbridge & Baker, 2020). We re-scored the boundary transformation scores to −1 (“closer”), 0 (“same”, only for Experiment 2), and 1 (“farther”) and calculated the mean boundary transformation score per image. With these two measures we 1) ran the correlation between mean aperture ratings and the boundary transformation scores and 2) computed the average boundary rating across images in each categorical aperture rating bin (see Figure 1A-B).

**Figure 1.**
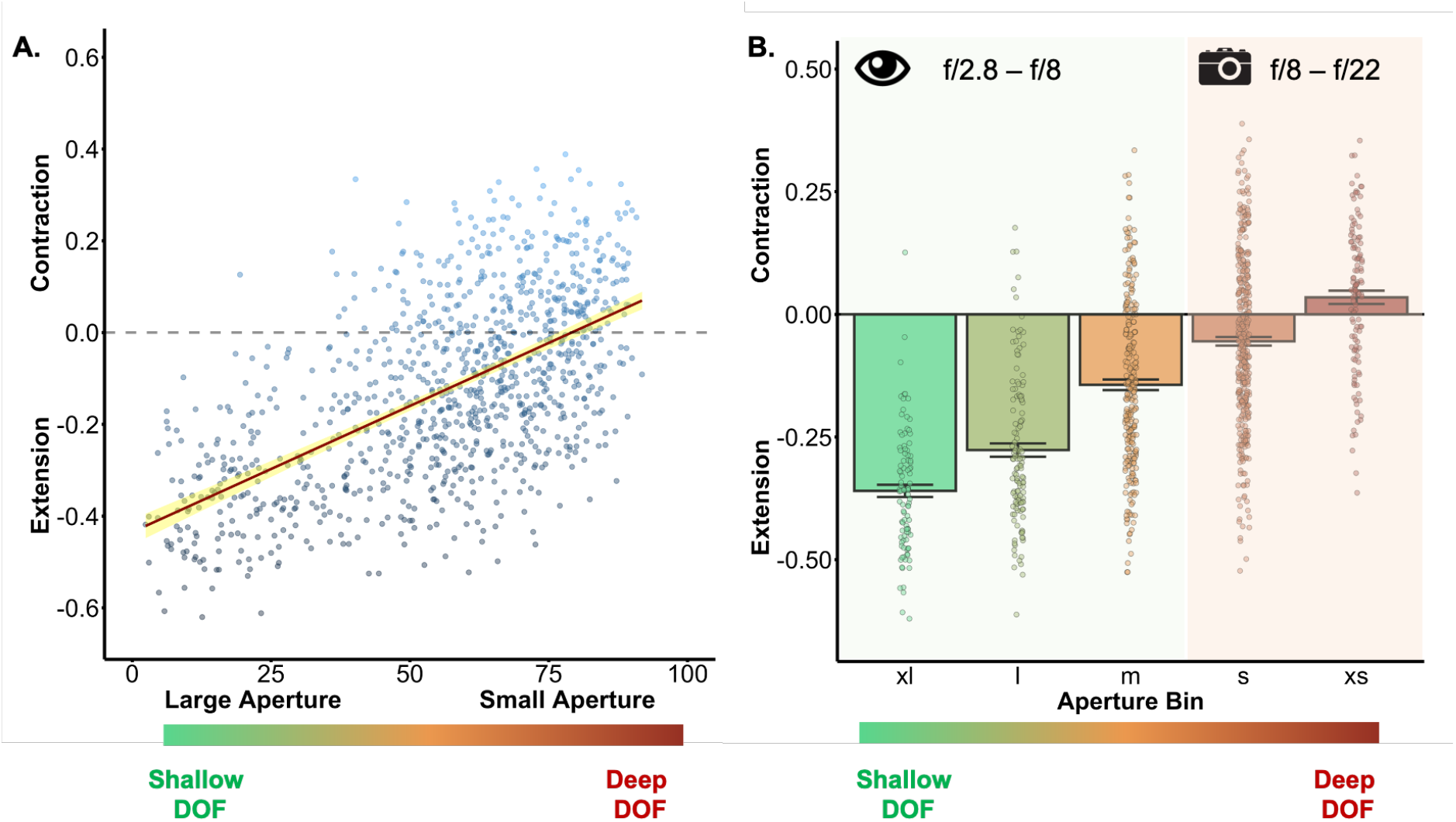
**(A)** Scatterplot with a linear regression line (in red), and 95% CI (in yellow). This shows the linear relation between aperture ratings and boundary transformation scores across the two 1000 images experiments of Bainbridge and Baker (2020). Images rated to be shot with large aperture led to strong boundary extension. Images rated to be shot with small aperture led to less boundary extension or even contraction. **(B)** Bar plot averaging boundary transformation scores across images for each of 5 aperture bins. Images in the S and XS bins were judged by professional photographers to be shot with apertures between f/8 and f/22, as computed from comparisons with aperture judgements of pictures with known aperture levels (Supplementary Methods). These apertures are mostly impossible for the human eye. Data points here represent mean boundary transformation value per image. Error bars represent the SE of the mean.

## Results

There was a strong and reliable correlation between rated aperture and boundary transformation score (Figure 1A, ρ = 0.58, *p* < 0.001). Images with a shallow DOF (large aperture) led to the largest extension effects, while images with deep DOF (small aperture) showed less extension or even contraction. To further inspect this relationship, we created 5 bins, each representing an interval of 20 on the aperture rating scale (XL – 0-20; L – 21-40; M – 41-60; S – 61-80; XS – 81-100). Compellingly, the bar plot in Figure 1B shows that contraction was reliable, on average, only for images rated to be shot with a small aperture (t-test vs 0 on the values of the XS bin - t (121) =2.56, *p* = 0.01, d = 0.23, 95% CI = [0.01, 0.06]). These aperture values are reachable by a camera but not by the human eye (Middleton, 1958; Hecht, 1987; see Supplementary Materials for how we estimated the *f*/stop value corresponding to each of the 5 bins shown in the figure).

Bainbridge & Baker (2020) showed that subjective ratings of viewing distance were correlated with boundary transformation scores, in line with other studies (Intraub et al., 1992; Hafri et al., 2021; Park et al., 2021). Images rated to have a close view led to larger extension than images rated to have a far-away view. It is possible that the relationship between rated DOF and boundary scores observed here is entirely absorbed by the relationship between subjective distance and boundary scores. To address this, we partialed out ratings of subjective distance. The correlation between aperture and boundary scores remained highly reliable (ρ = 0.29, *p* < 0.001), even when the shared variance between DOF ratings and subjective distance ratings was regressed out (Supplemental Materials for the same analyses splitting by image set, and partialing out other available image properties from Bainbridge & Baker (2020)).

### Experiment 2 and 3

In Experiment 1 we showed that boundary transformation scores and perceived DOF were related to each other across a large and diverse set of photographs, thus revealing that a previously unexplored image property affects memory for scene boundaries. However, as is the case for viewing distance (Bainbridge & Baker, 2020), the DOF of an image may covary with numerous other visual and semantic properties. For example, images with deep DOF are more likely to show an outdoor environment. The correlational approach used in Experiment 1 is thus unable to unambiguously ascribe the effects to DOF (or, indeed, viewing distance). Therefore, to firmly establish an effect of DOF on BE, we need to manipulate DOF while keeping all other properties equal. To do so, we shot the same 32 pictures of outdoor environments with two different camera apertures (f/5.6 and f/22). This factor was crossed with the camera’s focus distance, focusing either on a nearby or a faraway object in the scene (Figure 2). If the size of memory distortions depends on how the DOF of an image resembles natural viewing conditions, then images with shallow DOF (f/5.6) should show larger extension than images with deep DOF (f/22), independently of focus distance.

**Figure 2.**
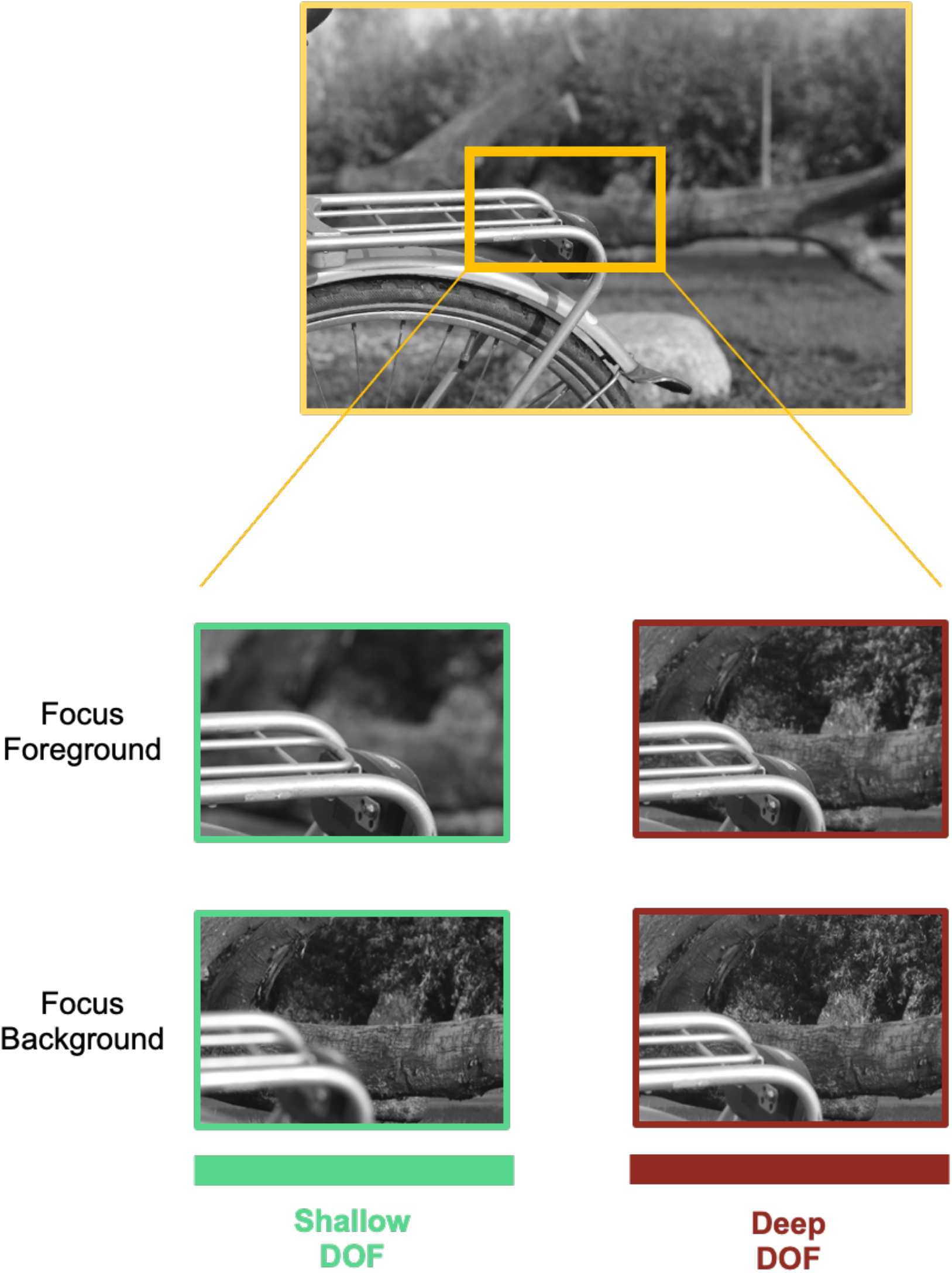
Example of the experimental conditions illustrated by a zoomed portion of one of the scene photographs. Each outdoor scene image was shot focusing either on a foreground object or on the background, at two levels of aperture – f/5.6 (shallow DOF) and f/22 (deep DOF). The full stimulus set is publicly available to download at the OSF repository, including the images equated for spatial frequency and histogram used in Experiment 3.

## Methods

### Participants

The sample size was determined through a power analysis based on the data of a pilot experiment (reported in Supplementary Materials). This analysis revealed that a sample size of 35 participants was required to obtain 80% power to detect a difference between large and small aperture conditions at least as large as the one observed in the pilot experiment, on the basis of a paired samples t contrast. In Experiment 2, thirty-nine participants were recruited for the experiment (16 females, mean age = 24.66 ± 4.10) to arrive at a final sample of N = 35 after exclusions based on missing data or low accuracy (see Analysis). In Experiment 3, thirty-six participants were recruited for the experiment (11 females, mean age = 24.36 ± 4.66). Due to server issues, out of 36 we obtained only 32 complete datafiles in Experiment 3 (see Analysis). Participants were recruited via Prolific (www.prolific.co) in return for monetary compensation at an hourly rate of 6.31 £/hr.

### Stimuli and apparatus

The stimulus set consisted of 6 photographs of 32 unique outdoor scenes (192 photographs). Each scene featured one or more objects in the foreground at a distance of approximately two meters, and one or more objects in the background at a distance of at least 5 meters. For each scene, 6 photographs were taken using a DSLR (digital single-lens reflex camera) with a 50mm fixed focal length lens placed on a tripod. Photos were shot at 3 aperture levels (*f/*5.6 (large); *f*/11.0 (medium); *f*/22 (small)), each one either manually placing focus on the foreground object(s) or on the background object(s) (See Figure 2). The medium aperture level was only used in the pilot experiment (see Supplementary Material). The ISO (light sensitivity of the sensor) was set manually in each scene (but kept constant across aperture and focus variations of that scene) and the shutter speed was automated to achieve similar lighting for the 6 photographs. We then generated “close view” scenes through resizing (without resampling) the original pictures. This was achieved by enlarging the image (i.e., zooming in) using a random percentage value between 17% to 24% (in steps of 0.5%) of the image’s surface area, and then cropping the image to its original size. Each scene was enlarged by the same percentage across the scene’s aperture and focus distance levels. All the pictures were then grayscaled and resized to 750 x 500 pixels. The images appeared on the screen at this resolution, and therefore varied in their size depending on the resolution the participants visualized them on their screen. Further, 8 different masks were generated by computing the average image of all the photos in the stimulus set and iterating image scrambling in 20×20 pixel blocks (randomly scrambling on the x and then on the y axis, or vice versa). For this and the following experiments, image processing was performed using the R package “Imager” (Barthelme & Tschumperle, 2019).

In Experiment 3, we matched the stimuli for low-level properties. Specifically, we used the SHINE toolbox (Willenbockel et al., 2010) running on Octave 5.1.0 (Eaton et al., 1997). Using the functions *sfMatch* and *HistMatch* we matched the spatial frequency and histograms of each scene across its focus and aperture levels. Therefore, for each scene four images were matched for their low-level properties – the small and large aperture images in both near and far focus levels. This procedure ensured that any difference between conditions in the experiment could not be due to covarying low-level properties emerging from the changes in focus distance and/or lens aperture. For both experiments, there were 128 images – 32 for each combination of aperture and focus distance. Further, we generated an additional 128 close view images belonging to the same conditions through resizing. Finally, the same 8 masks of Experiment 2 were used.

### Procedure

In each trial (Figure 3A), after a fixation cross with a random duration between 1 and 2 seconds, participants viewed a scene for 250 ms, followed by a 1s dynamic mask (changing every 250ms, 5 masks randomly selected from the 8 available masks in each trial), and then a closer or wider version of the same scene was shown again for 250 ms. Participants were asked to respond as fast and as accurately as possible pressing the “A” key if the second image looked closer or the “S” key if the image looked farther compared to the first image. Responses were given with the index (S) and middle (A) finger of the left hand. Right after response, or after 3.5 seconds, participants had to rate how sure they were of their response on a slider scale from 0 (“Not Sure”) to 100 (“Sure”). In the instructions, participants were told that the study concerned scene memory. Further, they were shown a self-paced example of the trial and performed a 10-trial practice block in which they were given feedback about their performance and got used to the task speed. Every 32 trials there was a self-paced break. The above procedure applies to both Experiments 2 and 3.

**Figure 3.**
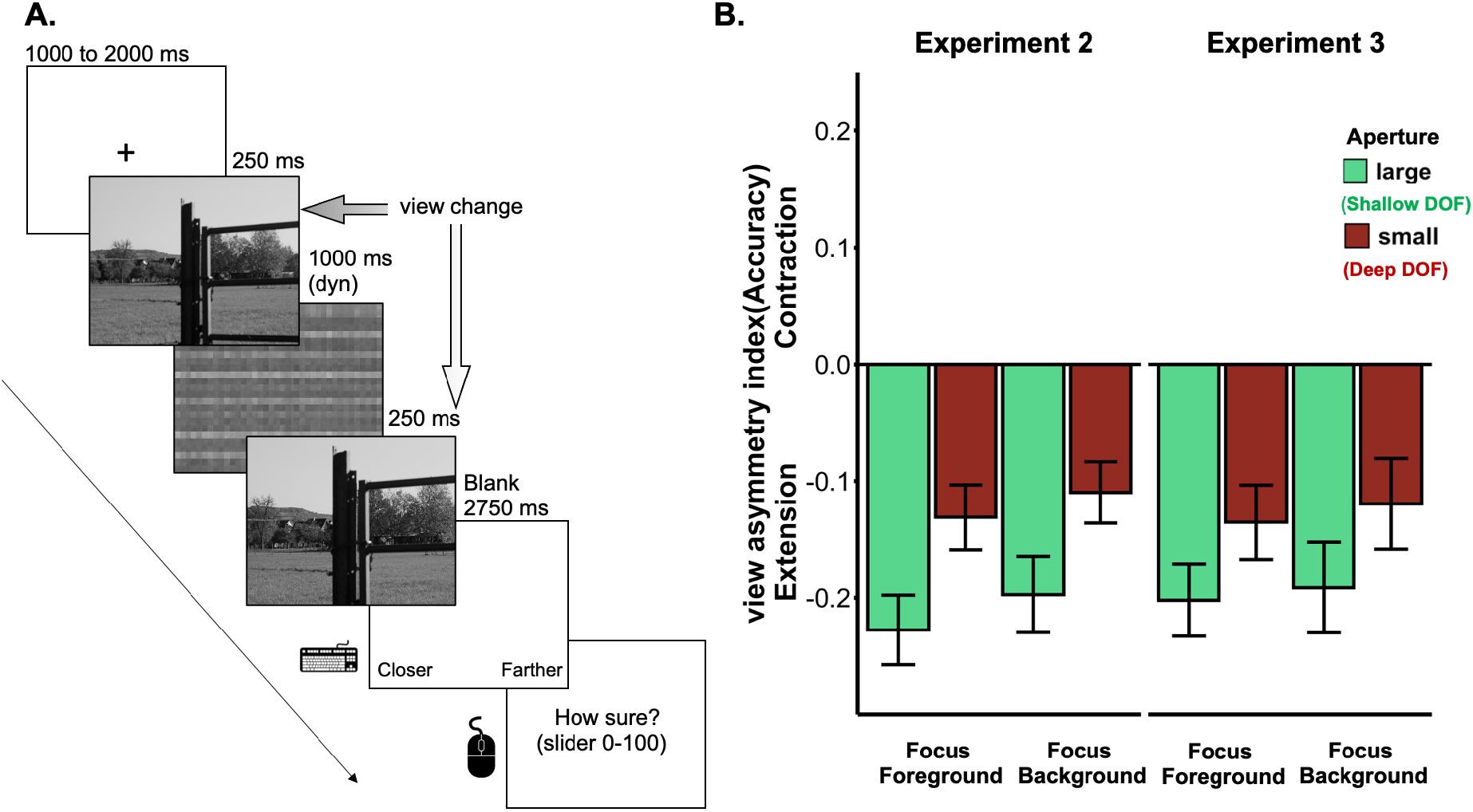
**(A)** Schematic representation of the procedure of Experiments 2, 3 and 4. The example shows a wide-to-close trial of Experiment 3. The probe picture shows a closer view than the first picture. **(B)** Results from Experiments 2 and 3. The View asymmetry index is computed as the difference in change detection accuracy for view changes (close-to-wide vs. wide-to-close). Negative values represent higher accuracy in detecting a change from wide to close, indicative of boundary extension. In both experiments there was an effect of apertures across both levels of focus distance. Error bars represent the SE of the mean.

### Design

The experiment consisted of 8 blocks of 32 trials. In each block, we ensured that the same scene was not presented more than once. In every block, there were 4 scene images for each of the experimental conditions (i.e., 2 views, close-to-wide/wide-to-close; 2 focus levels, far/near; 2 apertures, large/small). The full design thus included 256 trials in total.

### Analyses

In both experiments we excluded participants who did not have a complete datafile (1 participant for Experiment 2; 4 for Experiment 3) or who did not respond on more than 50% of trials (1 participant in Experiment 2; 0 in Experiment 3). Further, participants were considered outliers when their response time (RT) or accuracy was 2.5 SDs above (RT) or below (accuracy) the group mean across conditions (2 participants in Experiment 2; 0 in Experiment 3). The analyses for Experiment 2 thus included data from 35 participants, while those for Experiment 3 included data from 32 participants. Finally, for each experiment, we ran a 2 (View, close-to-wide/ wide-to-close view) x 2 (Aperture, large/small) x 2 (focus on foreground/background) repeated measures ANOVA on accuracy (correct detection of the second picture being closer or wider). For the purpose of visualization, we then computed a “view asymmetry index” subtracting accuracy on wide-to-close trials from accuracy on close-to-wide trials (Figure 3B). For this index, positive values indicate boundary contraction while negative values indicate boundary extension.

## Results

We used a paradigm that quantifies BE via a view-change detection asymmetry (Intraub et al., 1989; Intraub and Dickinson, 2008; Park et al., 2007; McDunn et al., 2014). In this paradigm (Figure 3A), all trials showed an actual change of view from the first to the second image, either changing from a wide to a close view or vice versa. Participants indicated whether the second image was closer or farther than the first image and rated their confidence on a slider scale. If the boundaries of the first image are extended in memory, participants should be worse in detecting changes from close to wide (vs. changes from wide to close) because the first (close) image is remembered to be wider than it was. Supplementary Materials report an experiment showing similar results when using the same paradigm used to obtain the BE scores employed in Experiment 1 (Bainbridge and Baker, 2020).

In both experiments, we found a main effect of view (Experiment 2: F(1,34) = 44.20, *p* < 0.001, η_p_^2^ = 0.57; Experiment 3: F(1,31) = 26.69, *p* < 0.001, η_p_^2^ =0.46). View changes from wide to close (Experiment 2: M = 0.80, SE = 0.02; Experiment 3: M = 0.81, SE = 0.02) were detected more accurately than view changes from close to wide (Experiment 2: M = 0.63, SE = 0.02; Experiment 3: M = 0.65, SE = 0.02). This effect indicates that boundaries were extended in memory when averaging across conditions.

Importantly, in both experiments we also found a significant View x Aperture interaction (Figure 3B, Experiment 2: F (1, 34) = 35.67, *p* < 0.001, η_p_^2^ = 0.51; Experiment 3: F(1,31) = 11.46, *p* = 0.002, η_p_^2^ = 0.27). Photographs with large aperture led to larger BE (Experiment 2: M = −0.21, SE = 0.3, t(34) = −7.75, *p* < 0.001, d = −1.31, 95% CI = [−0.27, −0.16]; Experiment 3: M = −0.20, SE = 0.3, t(31) = −6.17, *p* < 0.001, d = −1.09, 95% CI = [−0.26, −0.13])] than pictures with small apertures (Experiment 2: M = −0.12, SE = 0.02, t(34) = −4.83, *p* < 0.001, d = −0.82, 95% CI = [−0.17, −0.07]; Experiment 3: M = −0.13, SE = 0.03, t(31) = −3.74, *p* < 0.001, d = −0.66, 95% CI = [−0.20, −0.06]). View change did not interact with focus distance (Figure 3B, Experiment 2: F(1,34) = 2.24, *p* = 0.14, η_p_^2^ = 0.06; Experiment 3: F(1,31) = 0.48, *p* = 0.49, η_p_^2^ = 0.01), and there was no three-way interaction (Experiment 2: F(1,34) = 0.05, *p* = 0.82, η_p_^2^ = 0.01; Experiment 3: F(1,31) = 0.02, *p* = 0.88, η_p_^2^ = 0.01). The main effect of aperture (Experiment 2: F(1,34) = 4.12, *p* = 0.05, η_p_^2^ = 0.10; Experiment 3: F(1,31) = 0.04, *p* = 0.85, η_p_^2^ = 0.01), of focus distance (Experiment 2: F(1,34) = 0.23, *p* = 0.63, η_p_^2^ = 0.01; Experiment 3: F(1,31) = 0.22, *p* = 0.64, η_p_^2^ = 0.01), and the interaction between aperture and focus distance (Experiment 2: F(1,34) = 0.01, *p* = 0.96, η_p_^2^ = 0.01; Experiment 3: F(1,31) = 2.86, *p* = 0.10, η_p_^2^ = 0.08) did not reach significance. A similar pattern of results was observed for inverse efficiency scores (RT/% correct), confirming that results were not due to speed accuracy trade-offs (Supplementary Materials). Finally, the confidence ratings mirrored the accuracy results (Supplementary Materials).

### Experiment 4

In Experiments 2 and 3 we observed extension at all aperture levels, even when the DOF of the images was deep (and therefore not resembling human vision) and the focus was on the background (for which normalization may be expected). This could reflect the relatively close-up views of these photographs, with the background generally being within 50 meters of the camera. To test whether DOF also influences BE for very distant views of scenes (for which normalization accounts predict boundary contraction; Bainbridge & Baker, 2020), in Experiment 4 we used a new set of photographs depicting distant views of scenes, again with deep or shallow DOF (large vs small apertures).

## Methods

### Participants

We tested 38 participants (25 females, 2 other, mean age = 28 ±5 years) to arrive at the desired sample size of N=35 (based on a power analysis; see Methods of Experiments 2 and 3). Participants were recruited via Prolific (www.prolific.co) in return for monetary compensation at an hourly rate of 6.31 £/hr.

### Stimuli and apparatus

We shot 2 photographs of 28 unique outdoor scenes (56 images). Each scene featured a landscape in the background, often containing a path or a house at a far-away distance (Figure 4A). For each scene, we shot two photographs using a DSLR camera with a 45-55 mm focal length lens placed on a tripod. Photos were shot at 2 aperture levels (*f*/5.6 (large); and *f*/22 (small)), each one placing focus on the foreground plane (Stimuli are available to download at the OSF repository). The ISO (light sensitivity of the sensor) was set manually in each scene (but was held constant across aperture levels) and the shutter speed was automated to achieve similar lighting for the 2 photographs.

**Figure 4.**
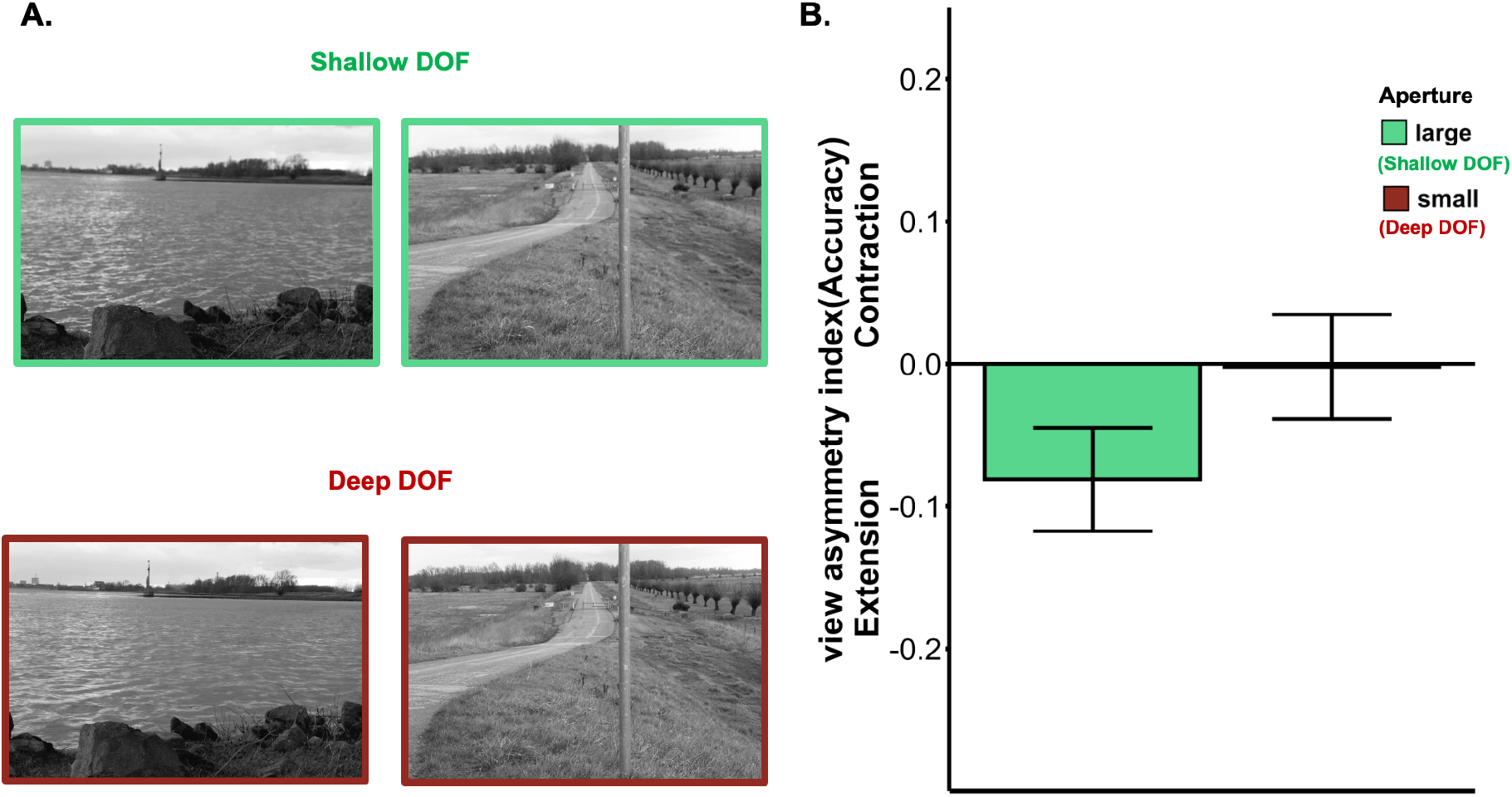
**(A)** Examples of two scenes from Experiment 4 with shallow and deep DOF. **(B)** Results from Experiment 4. The View asymmetry index is computed as the difference in change detection accuracy for view changes (close-to-wide vs. wide-to-close). Negative values represent higher accuracy in detecting a change from wide to close, indicative of boundary extension. This effect was present for the shallow DOF and absent for the deep DOF photographs. Conventions as in Figure 3.

All photographs were then gray-scaled and matched for luminance using the SHINE toolbox. Specifically, we matched the histogram of each pair of unique scene photographs across their aperture levels. We then resized the images to 750×500 pixels.

Finally, we generated “close view” scenes using the same procedure as in Experiments 2 and 3. Like before, the same scene was resized by an equal percentage across their aperture levels. In total, we had 112 images – 28 for each aperture level in their wide and close view. The masks were generated using the same procedure as used in Experiments 2 and 3. The size at which the images were presented was controlled by asking participants to place a real-world object of a standard size (a credit card) on the screen and use it to resize a rectangle displayed on it. Based on the ratio between the rectangle image width (in px) and the physical width of the card (in mm) we scaled the images so that they would appear 15 cm wide and 7.5 cm long on every screen, independently of screen resolution and size.

### Design

The experiment consisted of 2 blocks of 112 trials. Each block was further divided in 8 mini-blocks of 28 trials. Within each mini-block, the same scene was not presented more than once. In each mini-block, there were 7 scene images for each of the experimental conditions (i.e., 2 views, close-to-wide/wide-to-close; 2 apertures, large/small). The second 112-trials block had the same structure as the first block but the order of the mini-blocks was reshuffled. The full design included 224 trials in total.

### Procedure

The procedure was the same as Experiments 2 and 3.

### Analyses

The same exclusion criteria as Experiment 2 and 3 were used. Based on these, two participants were excluded because they did not respond to more than 50% of the trials and one participant was excluded because his/her accuracy was 2.5 SDs below the group mean across conditions. The analyses included data from 35 participants. As main analysis, we ran a 2 (View, close-to-wide/wide-to-close view) x 2 (Aperture, large/small) repeated measures ANOVA on accuracy.

## Results

We found a significant View x Aperture interaction (F(1,34) = 19.73, *p* < 0.001, η_p_^2^ = 0.37; Figure 4b). BE was reliable for the large aperture condition (M = −0.08, SE = 0.04; t(34) = −2.23, *p* = 0.03, d = −0.38, 95% CI = [−0.15, −0.01]) but was no longer observed for the small aperture condition (M = −0.002, SE = 0.04, t(34) = −0.06, *p* = 0.96, d = −0.01, 95% CI = [−0.08, 0.07], BF_10_ = 0.18). We did not observe a main effect of Aperture (F(1,34) = 1.42, *p* = 0.24, η_p_^2^= 0.04), nor a main effect of view-change (F(1,34) = 1.38, *p* = 0.25, η_p_^2^ = 0.04). A similar pattern was observed for inverse efficiency scores (RT/% correct), confirming that results were not due to speed accuracy trade-offs, and for confidence ratings (Supplementary Materials).

These results generalize the effects of DOF on BE to a new stimulus set consisting of distant scene views.

## General Discussion

In four experiments, we found that BE depends on a photograph’s DOF. In Experiment 1, across a large and variable image set, DOF was a strong predictor of BE, with BE being largest for images with shallow DOF, resembling human vision. By contrast, BC was only reliably observed for images with deep, unnaturalistic DOF. Three controlled experiments showed that DOF modulated BE even when keeping other properties of the scene constant across conditions. Altogether, these results demonstrate that BE is reliably observed for ecologically-representative stimuli.

Because DOF may covary with perceived distance across photographs, we ensured that the relationship between DOF and BE observed here could not be explained by differences in perceived distance between shallow and deep DOF. First, in Experiment 1, the relationship between DOF and BE remained reliable after regressing out subjective distance ratings. Second, in Experiments 2 and 3, focus distance did not interact with DOF, even though for large aperture focus distance was clearly visible (Figure 2). It should be noted, however, that previous work showed that BE is larger for scenes that show closer compared to far-away views (Intraub et al., 1992; Intraub & Richardson, 2008; Bertamini et al., 2005; Bainbridge & Baker, 2020; Park et al., 2021; Hafri et al., 2021). These results suggest an additional role for normalization processes. This more generic normalization process (also observed for objects) could exist independently of the more scene-specific predictive mechanism driving BE (Intraub et al., 1996; 1998). Experiment 4 supports this idea, showing no BE for distant scene views with deep DOF, suggesting that the opposite effects of normalization (leading to BC) and predictive (leading to BE) processes cancelled each other out for this condition. Importantly, predictive processes were strong for naturalistic (shallow) DOF, leading to BE even for distant scene views.

When integrating our results with previous findings, we see the largest BE for photographs with a shallow DOF, taken from relatively close distance, with the main object seen from a central vantage point (Gagnier et al., 2011). What do all these aspects have in common? One possibility is that images shot under these conditions best resemble how we naturally sample the visual world. Indeed, images with these characteristics show a more partial view of the scene than images with deep DOF, wide view and lateral vantage point. These partial views are more likely to require integration of visual input: When the image is less complete, observers may need to rely more on top-down expectations of scene layout, drawing on sources other than the visual input to complete the percept.

Another aspect that photographs with deep DOF, shot from large distance, and from a lateral vantage point have in common is that they typically contain many visible objects. To perceptually encode and remember such scenes requires greater attentional resources. This may lead to a loss of peripheral image content, leading to boundary contraction. Furthermore, fewer resources will be available for predicting what is beyond the scene’s boundaries, thus reducing BE. Nevertheless, our results suggest that the DOF effects on BE observed here cannot be fully accounted for by image complexity. First, the relationship between DOF and boundary scores remained reliable even when regressing out the number of objects in the images (Supplemental Materials). Second, in Experiments 2 and 3 DOF affected BE regardless of whether the camera’s focus was on the background (showing a larger number of objects) or on the foreground.

Based on our results, we propose that BE reflects a constructive mechanism with adaptive value that is conditional to a scene being perceived as naturalistic. An image shown at a plausible distance from the observer, with a DOF resembling the day-to-day perceptual experience, will likely lead to extrapolation of scene layout from memory, and therefore BE. Where one or multiple of these properties are removed then BE is attenuated. Future research is needed to identify those conditions of the external visual input that allow the mental representation of scenes, stored in memory, to complete external percepts.

By showing stronger BE for images with more naturalistic DOF, our results support the view that BE is a scene perception phenomenon rather than a phenomenon specific to photographs (Intraub, 2012). This is in line with previous evidence showing that BE is independent of photographical artifacts (e.g., image magnification; Bertamini et al., 2005), and that it occurs across modality (Intraub et al., 2015). The viewing conditions in our experiments differed substantially from our real-world visual experience. Our stimuli were brief static views, did not cover the full visual field, lacked binocular depth cues, and focus distance was not yoked to the observer’s fixation in the scene. Future studies are needed to test BE for more naturalistic viewing conditions, for example using virtual reality (Snow & Culham, 2021). We predict that BE will be reliably observed under those conditions.

Finally, our results have broad implications for disciplines aiming to study natural vision. Many recent psychological, neuroscientific, and computer vision studies implement large stimulus sets to achieve higher ecological validity (e.g., Hebart et al., 2020; Bainbridge & Baker, 2020; Chang et al., 2020; Mehrer et al., 2021). Our findings indicate that such large image sets are not necessarily representative of how observers perceive their visuo-spatial world. Indeed, there is evidence that decreasing a scene’s realism impacts visual memory (Tatler and Melcher, 2007). By showing that DOF can drastically change the influence of top-down knowledge on visual processing, our results imply that it is important to use images with naturalistic DOF in future work.

## Acknowledgments

We thank Surya Gayet, Simen Hagen, Genevieve Quek, and Lu-Chun Yeh for feedback on earlier versions of the manuscript, and the other Peelen Lab members for helpful discussions and feedback during lab meetings.

## Open data

All experiment scripts, raw data, analysis scripts, and stimuli for Experiment 2, 3 and 4 have been made accessible here (via the Open Science Framework): *https://osf.io/7btfq/*

## Author Contributions

M. Gandolfo performed data collection, visualization, analyses and programmed the experiments, M. Gandolfo and M.V. Peelen designed the experiments and wrote the manuscript, H. Nagele created stimuli for experiment 2 and 3, M. Gandolfo for experiment 4, M. Gandolfo and M.V. Peelen conceived the research.

