## Supplementary Materials for "Predictive processing of scene layout depends on naturalistic depth of field"

### Additional analyses – Experiment 1

*Partial correlations split by database*

Here we repeat the correlational analysis of Experiment 1, separately for the two image databases used in Bainbridge and Baker (2020), that is the GOI (Google Open Image database, Kuznetsova et al., 2020) and the SUN database (Xiao et al., 2016). The correlation between DOF ratings and boundary transformations scores was significant in both databases – Google database: $\rho = 0.53$, $p < 0.001$; Sun database $\rho = 0.23$, $p < 0.001$ (see **Figure S1ABC**). The same results held for the partial correlation, regressing out subjective distance ratings – Google database $\rho = 0.37$, $p < 0.001$; Sun database $\rho = 0.16$, $p < 0.001$.

*Partial correlations between aperture and boundary transformation ratings regressing out the number of objects*

We assessed whether the number of objects would explain the relationship between depth of field and boundary transformation scores by partialling out the number of objects of each image from the correlation between boundary transformation scores and depth of field ratings. The correlation remained of moderate size and highly reliable ($\rho = 0.54$, $p < 0.001$) even when regressing out the number of objects present in each scene.

*Partial correlations between aperture and boundary transformation ratings regressing out all the other variables*

In the following analysis we report the correlation between DOF ratings and boundary transformation scores after regressing out the contributions of all the other variables available for the images of Bainbridge and Baker (2020). These are: the number of objects in the image, the subjective distance, the object centricity (distance in pixels from the center), and the object size. The partial correlation between DOF and boundary transformation scores remained reliable ($\rho = 0.24$, $p < 0.001$) after regressing out all the other available ratings. This suggests that the relationship between depth of field and boundary transformation ratings cannot be fully explained by these other image properties.

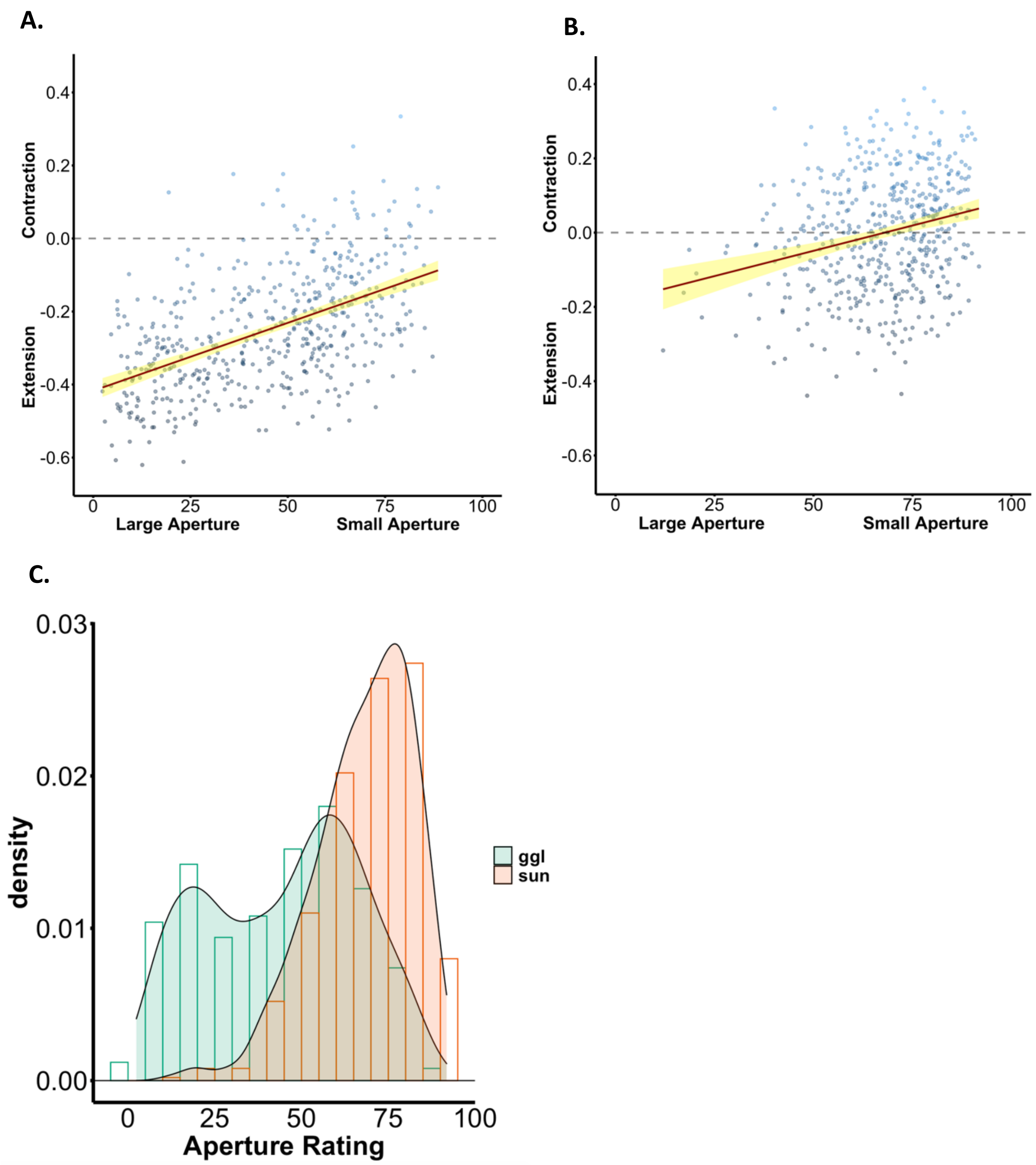


**Figure S1 –** Scatterplots illustrating the relationship between aperture ratings and boundary transformation scores separated by the two databases **A.** Google open images database and **B.** Sun database. **C.** Histogram illustrating the distribution of ratings between the Sun and the Google Open Image database.

### <u>Expert Photographers ratings</u>

The aim of this study was to estimate the f-stop aperture value (or range of values) to which the subjective rating bins of amateur photographers corresponded. To do so, we recruited professional photographers and asked them to judge a randomly selected subset of images belonging to each bin (XL – L – M – S – XS, see figure **S2** for an example of images coming from these bins) together with images with known aperture level.

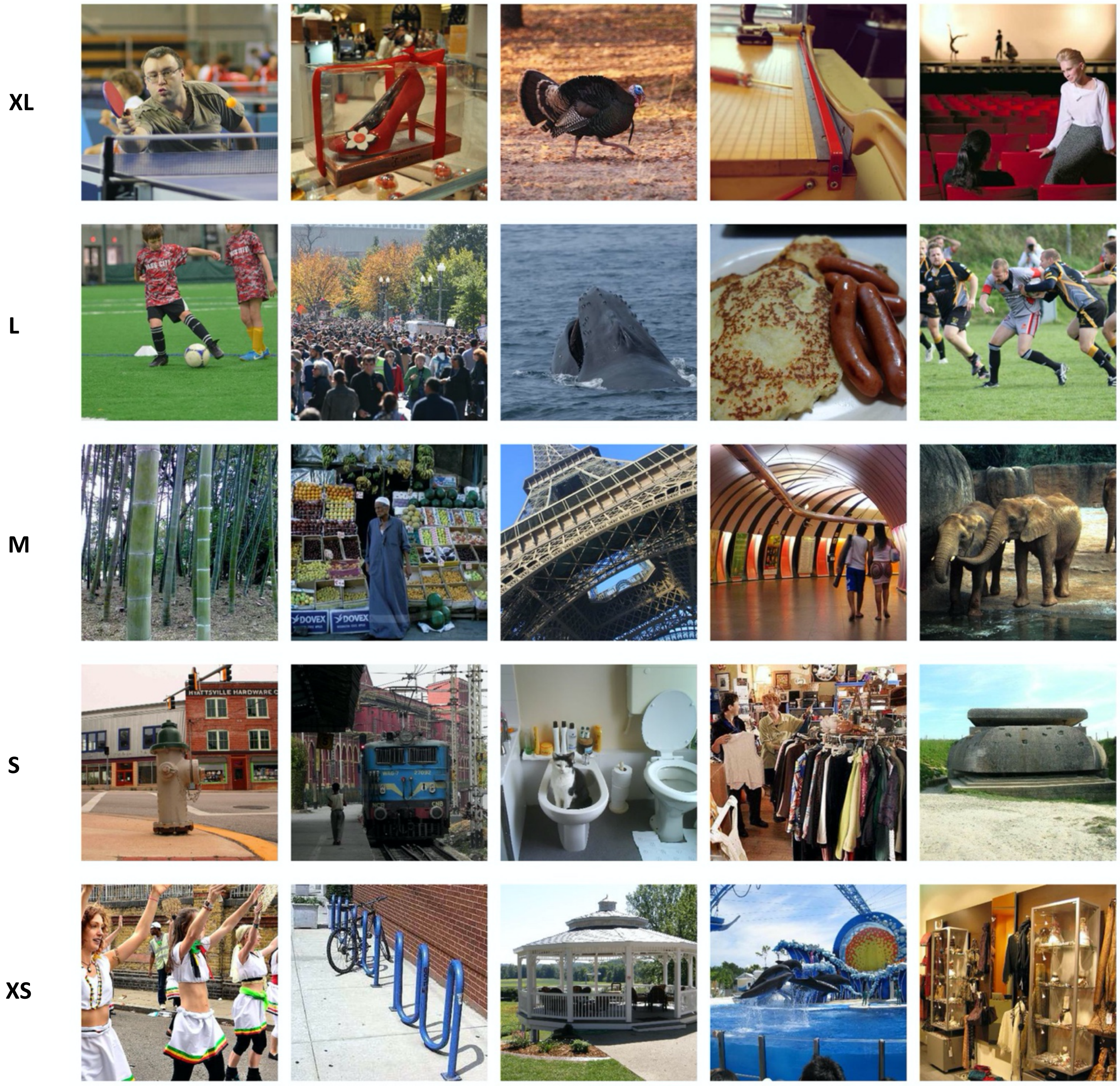


***Figure S2.*** *The image shows a sample of images from Bainbridge and Baker (2020) based on the aperture ratings bins from XL (shallow depth of field) to XS (deep depth of field).*

#### Participants

Participants were all self-proclaimed professional photographers recruited through social networks and snowball sampling. Sixteen photographers took part in this online study (9 males, 1 unknown, 6 females; mean age = 36 ± 9.85). Participation was on a voluntary basis.

#### Stimuli and apparatus

Based on the ratings of Experiment 1 we randomly selected 60 images with a mean rating falling within each of the aperture bins (0-20 (XL); 21-40 (L); 41-60 (M); 61-80 (S); 81-100 (XS)) for a total of a subset of 300 images from Bainbridge and Baker (2020). Further, we downloaded from www.shutterdial.com 105 images, 15 for each of 7 f-stop aperture values (f/2.8; f/4.0; f/5.6; f/8.0; f/11.0; f/16.0; f/22.0).

#### Design

Each participant completed 1 of 6 versions of the experiment**.** Each version included 50 images from Experiment 1, 10 for each of the 5 aperture bins. Further, 28 more images per participant were randomly selected from the pool of stimuli with known aperture level, 4 for each of the 7 *f*-stop values. The experiment included 78 trials in total.

#### Procedure

At the beginning of the experiment participants were told that they were recruited as professional photographers and that their expert eye was required to assess the f-stop value of images coming from the web. First, they completed the same questions of experiment 1A related to their experience and knowledge of photography concepts. Afterwards, participants were briefly shown an example image at different levels of aperture (therefore with shallower or deeper depth of field), participants were informed that they will have to guess the aperture level of images by choosing one out of seven f-stop options (f/2.8; f/4.0; f/5.6; f/8.0; f/11.0; f/16.0; f/22.0) using the mouse. When an image with known aperture level appeared (~ 1/3 of the total number of trials) participants received feedback about their performance. The feedback was adaptive, in that depending on the distance between the button pressed and the correct button a different message was displayed together with the actual f-stop level of the image. If the distance was 1, participant received: “Almost! The correct aperture was ..”; if distance was 2 “Not too far! The correct aperture was ..”; if it was 3 “Not quite! The correct aperture was .. ”; if distance was 4 or more “Wrong! The correct aperture was ..”. At the end of the experiment a link to a document explained the hypothesis in more detail, briefly illustrating the concept of the boundary extension illusion.

#### Analyses

In this experiment we aimed to estimate the f-stop values of images belonging to each of the aperture bins identified with the slider ratings of Experiment 1. To do so, we extracted the average button press for the images with known aperture and then estimated the f-values of the bins as the two neighboring f-values corresponding to the average button press of each bin (button press scores ranged from 0 – f/2.8 to 6 – f/22). Before performing the analyses, we excluded 4 photographers depending on their answers to the quiz at the beginning of the experiment, 3 of those declared to have only “some experience with SLR photography” (i.e. not extensive knowledge of manual mode) and 1 answered wrongly to more than 3/5 (5/5) multiple-choice questions of the quiz.

#### Results

The 5 bins corresponded to the following aperture values: XL= f/2.8-f/4.0; L= f/4.0-5.6; M= f/5.6-f/8.0; S= f/8.0-f/22; XS= f/8.0-f/22. Note that participants could not distinguish between f/11, f/16, and f/22, responding to these pictures with highly similar average button press values (4.1, 4, and 4, respectively)

### Pilot Experiment

In this experiment, we measured BE by asking participants whether a second image *looks* closer or farther than a previously observed image, following the paradigm of Bainbridge and Baker (2020). While this paradigm has proven effective in revealing BE, the ratings reflect a subjective impression rather than objective performance (Liu et al., 2016). In the main experiments reported in the paper, we used a paradigm that quantifies BE via a view-change detection asymmetry (Intraub et al., 1989; Intraub and Dickinson, 2008; Park et al., 2007; McDunn et al., 2014).

#### Methods

##### *Participants*

Seventy-eight participants signed up for the online experiment (65 females, mean age = 20.62 ± 4.19 years). Participants were students recruited via the University's participant panel (SONA) in return for course credits. Because we had no a-priori hypothesis on the size of the effect this sample size was determined by the student availability on the participant panel of the University in a period of four weeks.

##### *Stimuli and apparatus*

The same stimulus set of the experiments in the main text was used, now also including the medium (f/11.0) aperture level. For this experiment, we included only those images showing a "wide" view. Three scenes (two from the main experiment and 1 new scene), including photographs for each of the aperture and focus conditions, were used as catch trials. For these three photographs, we generated zoomed-in versions by enlarging the surface area of the images by 24% (in line with the procedure of Intraub and Dickinson (2008)) and then resizing (i.e., cropping) them to their original size.

##### *Procedure*

The experiment mimicked the procedures of the 1000-image RSVP Experiment 2 of Bainbridge and Baker (2020), a paradigm commonly used to measure BE effects (Park & Konkle, 2021, Intraub & Dickinson, 2008; Figure S3A). In each trial, participants saw a scene presented for 250 ms, followed by a 250 ms dynamic mask (5 masks randomly selected from the 8 available masks for each participant, presented for 50 ms each), and then the same scene was shown again. After 1 second, a response screen appeared for 3 seconds or until response. The response screen displayed text asking whether the second image was closer (scored as -1, "J" key), the same (scored as 0, "K" key) or farther (scored as +1, "L" key) than the first image.

In the instructions, participants were informed that the study concerned scene memory. They were also informed that the picture to be judged could have been either taken from the same, a closer-up, or a farther away view. Importantly, in the trials of interest the image to be judged was the same as the first image. In the catch trials (10% of the total number of the test trials), the second image was either an obviously zoomed-in or a zoomed-out version of the first photo. The catch-trials allowed to 1) have attentive participants to reinforce their belief that there were actual changes in the images and 2) have an accuracy measure that could be taken as an index of data quality (i.e., a suitable attention check for this task). Participants were invited to take a break at the end of each block (every 33 trials).

#### *Design*

The experiment consisted of 6 blocks of 33 trials. In each block, we ensured that the same scene was not presented more than once. In every block, there were 6 scene images for each of the experimental conditions (i.e., 2 focus levels, foreground/background x 3 apertures, large/medium/small). Further, there were three catch trials (with the second image being actually zoomed in or out). These were selected randomly for each participant from a pool of 36 possible trials (i.e., 2 focus levels x 3 apertures x 2 zoom types x 3 photos), making sure that no catch trial image was repeated within a block. The full design thus included 198 trials, 180 experimental trials and 18 catch trials.

#### *Analyses*

Based on the catch-trials performance we excluded participants who were below or at chance accuracy in the catch trials (mean accuracy <= 33%), the participants who responded more than 90% of trials “same” (excluding the catch trials), and the participants who did not respond (letting the trial to time out) in more than 50% of the total number of experimental trials. Based on these criteria we excluded 12 participants (6 for low-catch trial performance; 5 for not responding in more than 50% trials; 1 participant for responding “same” in > 90% of the experimental trials). The analyses thus included data from 66 participants. We then scored “closer” responses to -1, “same” responses to 0 and “farther” responses to 1. Finally, we computed the mean responses per participant in each condition and ran a 3 (aperture levels) x 2 (focus on foreground/background) repeated measures ANOVA.

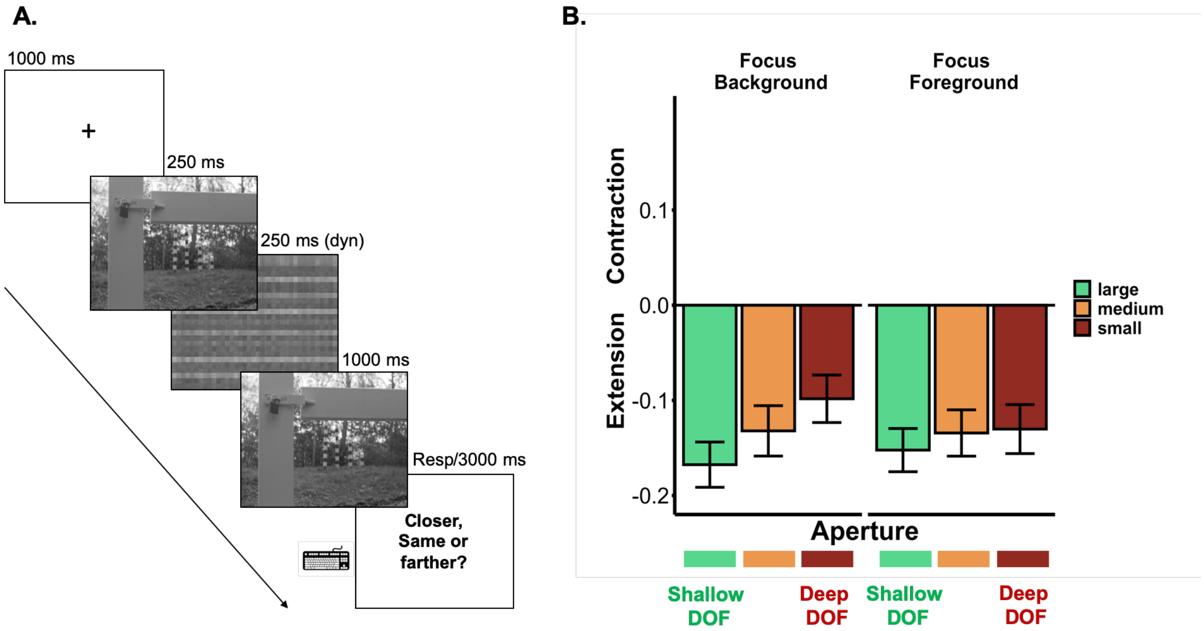


**Figure S3. (A)** Schematic representation of the pilot experiment. The procedure matched Bainbridge and Baker’s (2020) 1000 images Experiment 2, thus including the “same” response option. **(B)** Bar plot showing the main effect of aperture, F(2,130) = 5.27, *p* = 0.006, occurring across both focus distance levels. Images with large aperture led to the largest extension. Error bars represent the SE of the mean.

**Results**

Confirming our hypothesis, we found a main effect of lens aperture, $F(2, 130) = 5.27$, $p = 0.006$, $\eta_p^2 = 0.08$), showing stronger BE (i.e., second image rated as closer than the first for "same" trials) for images shot with large aperture, f/5.6 ($M = -0.16$, $SD = 0.18$) than images shot with medium aperture, f/11 ($M = -0.13$, $SD = 0.19$; $t(65) = -2.31$, $p= 0.02$, $d = -0.28$, 95% CI = [-0.53, -0.04]) and images shot with small aperture, f/22 ($M = -0.11$, $SD = 0.18$; $t(65) = -2.79$, $p= 0.007$, $d = -0.34$, 95% CI = [-0.59, -0.09]). There was no significant difference between medium and small aperture stop ($t(65) = -1.27$, $p= 0.21$, $d = -0.16$, 95% CI = [-0.05, 0.01]). Thus, while boundary extension was found in all conditions, it was reduced when images showed deeper DOF (Figure S3B). The effect of aperture did not interact with focus distance ($F(2, 130) = 0.64$, $p = 0.53$) and there was no main effect of focus distance ($F(1,65) = 0.99$, $p = 0.32$).

**Supplementary Analyses: Experiment 2 and 3**

**Inverse Efficiency Scores**

For Experiment 2, the ANOVA on the inverse efficiency scores (RT/mean accuracy) showed a main effect of View ($F(1,34) = 32.77$, $p < 0.001$, $\eta_p^2 = 0.49$). Participants detected changes from a wide to a close view ($M = 1069$, $SE = 55$) more efficiently than changes from a close-to-wide view ($M = 1610$, $SE = 91$). Crucially, similarly to what we report in the analyses on the proportion of correct responses, we also found a significant View x Aperture interaction ($F(1,34) = 13.12$, $p < 0.001$, $\eta_p^2= 0.28$). The asymmetrical performance in view detection indicated larger BE for pictures shot with large apertures ($M = -656$, $SE = 107$) than pictures shot with small aperture ($M = -425$, $SE = 91$).

In Experiment 3, the ANOVA on the inverse efficiency scores showed a main effect of View ($F(1,31) = 20.76$, $p < 0.001$, $\eta_p^2 = 0.40$). Participants detected changes from a wide to a close view ($M = 1089$, $SE = 97$) more efficiently than the converse ($M = 1567$, $SE = 79$). We also found the same View x Aperture interaction ($F(1,31) = 12.63$, $p = 0.001$, $\eta_p^2 = 0.29$). The asymmetrical performance in view-change detection indicated larger BE for pictures shot with large aperture ($M = -589.25$, $SE = 118$) than for pictures shot with small aperture ($M = -366$, $SE = 99$).

**Confidence Ratings**

In Experiment 2 we found a main effect of View ($F(1,34) = 42.12$, $p < 0.001$, $\eta_p^2 = 0.55$). Participants were more confident about their responses when judging changes from wide-to-close ($M = 67.57$, $SE = 1.80$) than changes of view from close-to-wide ($M = 59.77$, $SE = 2.15$). No other effect reached significance (all other $p$s > 0.15).

In Experiment 3 we also found a main effect of View ($F(1,31) = 14.50$, $p < 0.001$, $\eta_p^2 = 0.33$). Participants were more confident about their responses when judging changes of view going closer ($M = 69.63$, $SE = 1.84$) than changes going farther ($M = 63.13$, $SE = 1.94$). Further, for this experiment, we also found a View x Aperture interaction ($F(1,31) = 6.69$, $p = 0.01$, $\eta_p^2 = 0.18$). In line with the performance results, the asymmetry between wide-to-close and close-to-wide views was reliably larger for pictures shot with large aperture (M = -7.74, SE = 1.75) than for pictures shot with small aperture (M = -5.25, SE = 1.80).

### Supplementary Analyses: Experiment 4

#### Inverse Efficiency scores

In Experiment 4, the ANOVA on the inverse efficiency scores (RT/mean accuracy), similarly to what we report in the analyses on the proportion of correct responses, showed a significant View x Aperture interaction (F (1,34) = 9.21, *p* = 0.005, $\eta_p^2$ = 0.21). Importantly, when pictures were shot with large aperture, efficiency was higher for wide-to-close (M = 1294.16, SE = 70.34) than close-to-wide (M = 1562.49, SE = 96.76) changes (t (34) = -2.53, p = 0.016, 95% CI = [-484.02, -52.64], d = -0.43). Mirroring the accuracy results, this difference was absent (close-to-wide – M = 1493.09, SE = 89.24; wide to close – M = 1474.12, SE = 119.29) for the images shot with small aperture (t (34) = -0.13, p = 0.89, 95% CI = [-307.06, 269.14], d = -0.02, BF10 = 0.18). No other effects reached significance (all *p*s > 0.18).

#### Confidence

The following analysis included 34 participants. This is because one participant did not respond to the confidence slider and let it timeout for all but two trials. We found a main effect of View (F(1,33) = 12.62, *p* = 0.001, $\eta_p^2$ = 0.28). Participants were more confident about their responses when judging changes of view going closer (M = 61.71, SE = 2.47) than changes going farther (M = 59.68, SE = 2.34). Further, the View x Aperture interaction was significant, showing that the confidence results went in the same direction as the performance results (F (1,33) = 4.84, *p* = 0.035, $\eta_p^2$ = 0.13). When pictures were shot with large aperture, rated confidence was higher for wide-to-close (M = 62.82, SE = 2.51) than close-to-wide (M = 58.37, SE = 2.32) changes (t (33) = -4.44, p < 0.001, 95% CI = [-7.20, -2.68], d = -0.76). Mirroring the accuracy results, this difference was absent for the images shot with small aperture (t (33) = -1.47, p = 0.15, 95% CI = [-4.41, 0.70], d = -0.25, BF10 = 0.49). The main effect of aperture did not reach significance (F(1,33) = 2.45, *p* = 0.13, $\eta_p^2$ = 0.07).
